# Is torpor a quiescent state? Periodic motility and transient brain activation during daily torpor in Djungarian hamsters

**DOI:** 10.64898/2026.01.25.701526

**Authors:** Natalie L. Hauglund, Ritika Mukherji, Xiao Zhou, Anna Hoerder-Suabedissen, Rong Mao, Stuart N. Peirson, Annika Herwig, Tom Deboer, Vladyslav V. Vyazovskiy

## Abstract

Torpor is a hypometabolic state employed by many mammalian and non-mammalian species to cope with harsh environments. When exposed to a short photoperiod, Djungarian hamsters (*Phodopus sungorus*) enter daily torpor with body temperatures dropping to as low as 15°C. Despite the widely-held notion that torpor is a form of ‘deep sleep’, torpid animals are not completely inactive but exhibit occasional movements reflected in an increase in EMG tone. Little is known about these ‘EMG events’ during torpor and whether they have a functional role during the torpid state. We here analysed EEG, EMG, and brain temperature data from Djungarian hamsters, and used an automatic detection algorithm to identify periods of EMG activation during spontaneous daily torpor. The hamsters exhibited regular periods of motility that were invariably initiated during a decline in brain temperature and were followed by a brain temperature increase. The frequency of EMG events exhibited a negative correlation with brain temperature, such that lower brain temperature was associated with a higher frequency of EMG events. In addition, EMG events were associated with a pronounced increase in EEG power, especially between 9.5-15.5 Hz, which often started with an EEG pattern similar to an evoked potential preceding the increase in the EMG activity. On the contrary, micro-arousals during normothermic NREM sleep were associated with a decrease in EEG power, a decrease in brain temperature and were of shorter duration than torpor EMG events, indicating that the two phenomena may serve different purposes. We speculate that periodic motility associated with increased brain activity during torpor may play a role in thermoregulation, and help retain vigilance to potentially mitigate predation risk during this hypometabolic state.

## Introduction

Torpor is a ubiquitous, naturally occurring energy-conservation strategy characterised by hypometabolism and hypothermia(1–4). Some animals can enter states of hypometabolism in response to or in anticipation of harsh environmental conditions, such as extreme winters in Siberian regions(1–3), or hot and arid conditions of tropical deserts(4), where food(5) and/or water availability(6) is sparse. The onset of the torpor season is regulated mainly by a shortening photoperiod, and, together with adaptations in body mass, the reproductive system and fur changes, enables animals to lower their daily energy expenditure by more than 30%. Hence, spontaneous daily torpor is of special interest due to its translational potential for understanding controlled hypothermia and hypometabolism in a clinical setting and for enabling survival in extreme conditions, such as long-duration space flight(7–9).

The Djungarian hamster (*Phodopus sungorus*) undergoes hours-long periods of daily torpor during the winter months when kept under natural temperature and photoperiod conditions (10). This adaptive response can be reproduced in a laboratory setting by shortening the photoperiod, which causes hamsters to undergo gradual and easily trackable phenotypic modifications over 10-16 weeks, including reductions in body mass and changes in pelage. These changes reflect underlying physiological adaptations as the photoperiod shifts from summer-like long days to winter-like shorter days(2, 11–13), and can be used to predict the approximate week or month during which an animal is most likely to enter torpor(14).

Animals enter daily torpor via non-rapid eye movement (NREM) sleep(15–18). During the transition phase, there is a drastic reduction in rapid eye movement (REM) sleep, likely since REM sleep is associated with a loss of thermoregulatory control. Unlike preceding NREM sleep, during which animals are largely immobile and resting in a characteristic posture, the transition phase from NREM sleep to torpor is dynamic with periods of hyperventilation and/or shivering, as well as movements accompanied with changes in metabolic rate and body temperature. Once the lowest body temperature and metabolic rate are reached, animals typically maintain this state for several hours before initiating arousal from torpor. During body temperature decline and maintenance of hypothermia, the EEG activity lowers in amplitude but retains features resembling NREM sleep, such as EEG slow waves(19).

Similar to NREM sleep, periods of immobility during torpor are interspersed with occasional desynchronised higher frequency EEG with increased EMG activity, which are generally interpreted as waking bouts(16, 19). Although these arousal-like periods during torpor have been noted previously, their physiological relevance remains unclear. This highlights that naturalistic torpor is not a homogenous state of inactivity or immobility – an important feature that differentiates it from several other synthetically induced states of hypometabolism. In addition, torpor is known to be followed by a rebound of slow wave activity (SWA), which clearly distinguishes it from NREM sleep(14, 25). Thus, a deeper exploration of the neural dynamics of torpor is warranted.

In this study, we conducted a detailed analysis of a previously published dataset of EEG, EMG and cortical temperature during sleep and torpor in hamsters to gain a better understanding of the role and implications of brief periods of muscle activity (EMG events) occurring during torpor. The analysis revealed that torpor EMG events are characterised by dynamic changes in EEG and brain temperature, and that lower brain temperature is predictive of a higher number of torpor EMG events. In addition, we found that EMG events during torpor have different characteristics than micro-arousals during NREM sleep, suggesting that torpor EMG events have functions specific to the torpid state. Complementary experiments with detailed video recordings of hamsters during torpor further revealed that it is a highly dynamic state of vigilance, far from being a state of complete quiescence or lethargy, as it is often conceptualised.

## Results

### Physiological parameters of torpor

Adult, male hamsters (n=7) implanted with EEG electrodes, EMG electrodes, and a thermistor over the frontal cortex were recorded chronically for several weeks. Every animal exhibited bouts of torpor, which typically lasted for several hours (figure 1). One torpor bout from each hamster was used for analysis. During the analysed torpor bouts, hamsters on average spent 4.84 ± 0.71 hours in deep torpor (defined as brain temperatures being within 10% of the lowest torpor-associated brain temperature). Torpor bouts were associated with a decrease in EEG amplitude (measured over the frontal cortex) and a pronounced drop in brain temperature as described previously(2, 15, 16, 25, 26) (figure 1A-B). During torpor nadir, the brain temperature was on average 14.6°C lower than during euthermic NREM sleep (35.0±0.7°C in NREM sleep and 20.4±1.4°C in torpor) (figure 1C). During days without torpor, euthermic hamsters cycled through wakefulness, NREM sleep and REM sleep. Consistent with prior observations, analysis of the EEG activity during each state showed that wakefulness, NREM sleep and REM sleep exhibited markedly higher EEG power compared to torpor (figure 1D-F).

**Figure 1.**
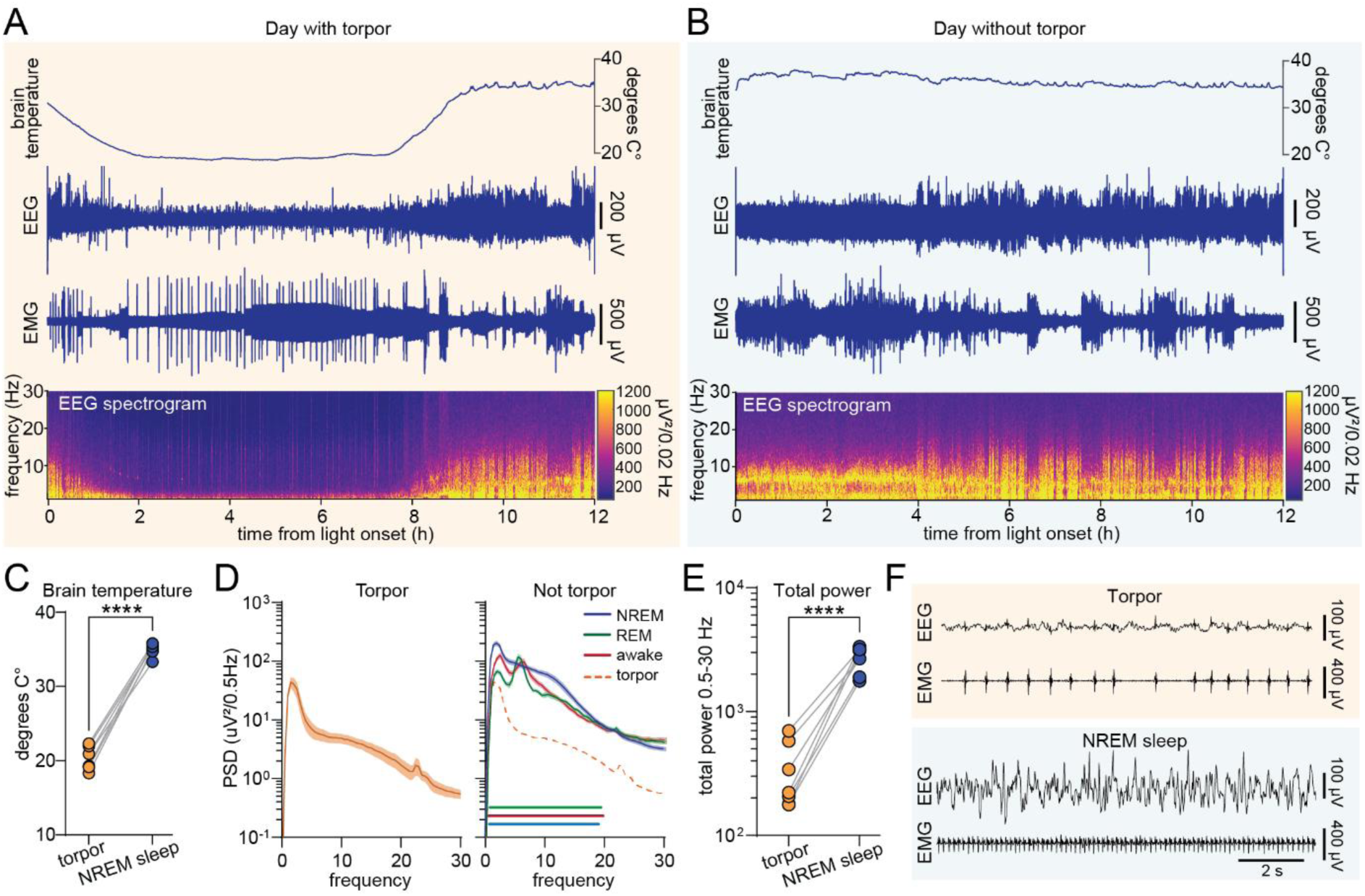
Physiological parameters of torpor. (A-B) Example of brain temperature measured above the frontal cortex (top), EEG, and EMG (middle), and EEG spectrogram (bottom) from the same hamster during a day with torpor (A) and a day without torpor (B). (C) Brain temperature during torpor and NREM sleep (n=7, paired t-test, P<0.0001). (D) Power spectral density plots of the EEG signal during torpor, wakefulness, NREM sleep and REM sleep (n=7, RM 2-way ANOVA with Sidak’s multiple comparison, dotted orange line depicts the average torpor PSD, horizontal lines denote frequencies with significant difference between torpor compared to wakefulness, NREM and REM sleep). (E) Sum of EEG power between 0.5-30 Hz during torpor and NREM sleep (n=7, paired t-test, P<0.0001). (F) Example of EEG and EMG signals from a hamster during torpor (top) and NREM sleep (bottom). Error bars are ±SEM. PSD = power spectral density.

### Low cortical temperature is associated with higher EMG event frequency and duration

Along with the diminished EEG amplitude and decreased temperature, torpor bouts were characterized by frequent events of muscle activity (figure 2A). These EMG events were associated with an increase in brain temperature that on average began ∼30 seconds after the beginning of the muscle activity (figure 2B). To analyse potential variance in brain temperature, EMG event duration, and EMG event frequency over the course of the torpor bout, the full period comprising the torpor nadir (defined as the brain temperature being within 10% of the lowest recorded brain temperature) was divided into five consecutive time bins. The durations of the time bins were dependent on the duration of the torpor bout and therefore differed between animals but were on average 58.18 ± 8.46 minutes long. Analysis of the brain temperature in each time bin showed that the lowest brain temperatures were detected around the middle of the torpor bout (20.0 ± 1.4°C) (figure 2C). Analysis of the duration of all EMG events across the torpor nadir revealed a bimodal distribution with the majority of EMG events lasting approximately 30-36 seconds and a smaller subset with a shorter duration below 12 seconds (figure 2D). Interestingly, EMG event duration and brain temperature over the course of the torpor bout showed an inverse pattern with longer duration EMG events in the middle of the torpor bout when brain temperatures were lowest (figure 2E). While the duration of the inter-EMG event interval (time between two consecutive events) did not vary significantly as a function of time within the torpor bout (figure 2F), the incidence of EMG events increased over the duration of the torpor bout and peaked with an average frequency of 8.0 ± 2.4 per hour thus occurring on average every ∼7.5 minutes (figure 2G). To explore the relationship between EMG event duration, EMG event frequency and brain temperature, the average value for each parameter was calculated for each of the five time bins spanning the torpor bout, such that each hamster contributed with five values to the correlation. There was no correlation between the average EMG event duration and the average EMG event incidence (figure 2H). Both EMG event frequency and EMG event duration exhibited a negative correlation with brain temperature during the EMG event (figure 2I-J). The strongest correlation was between EMG event frequency and brain temperature. Colour coding of data points from each hamster revealed that data points from the same hamster tended to form clusters, indicating that this correlation was mainly driven by differences in the absolute brain temperature and EMG event frequency across hamsters. Thus, hamsters with the lowest brain temperature generally had the highest frequency of EMG events.

**Figure 2.**
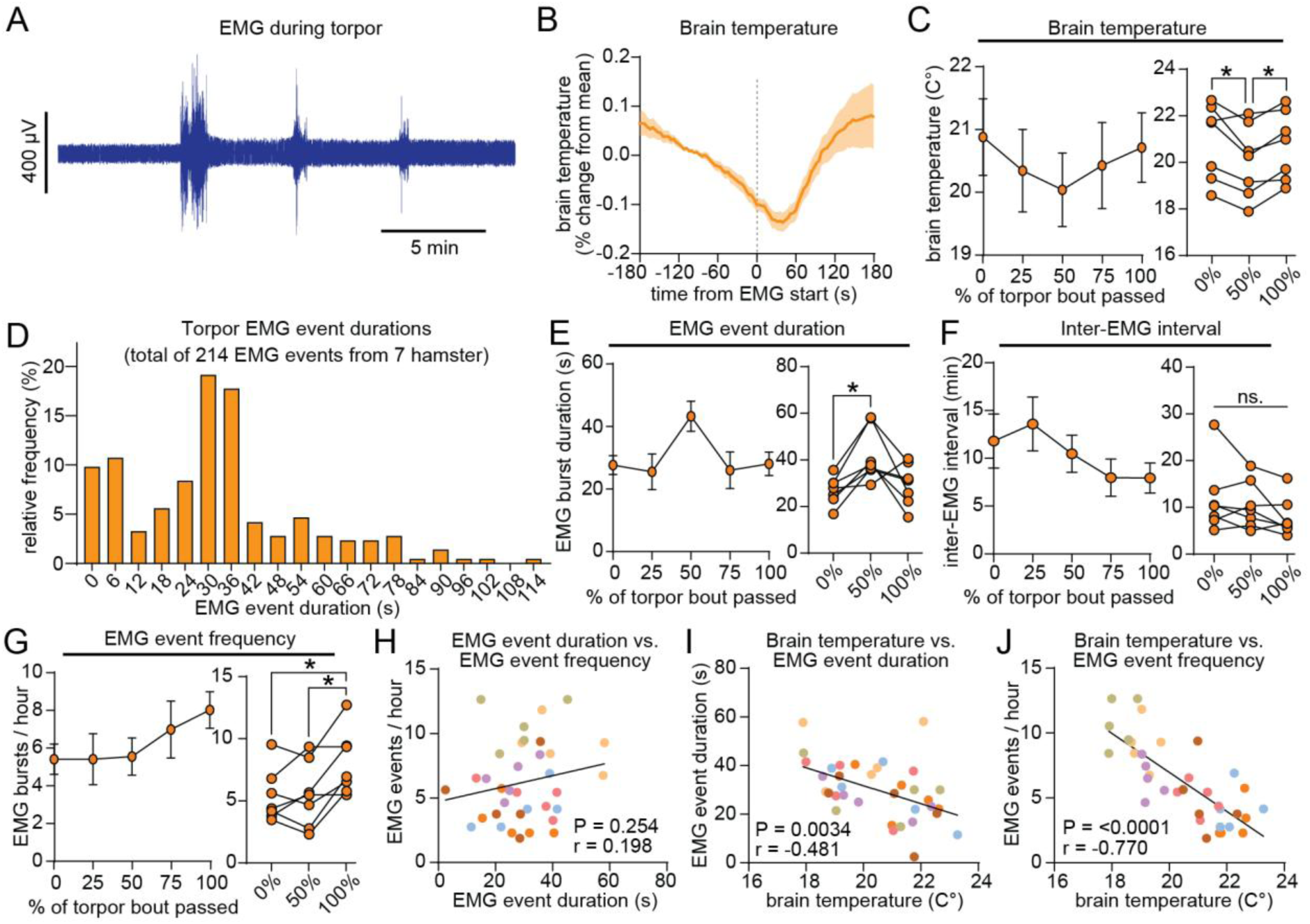
Low cortical temperature is associated with higher EMG event frequency and duration. (A) Example of EMG trace from a hamster during torpor showing periods of muscle activity. (B) Average brain temperature time-locked to muscle activity during torpor (n=7). (C) Average brain temperature during a whole torpor bout expressed as percent of torpor bout that has passed (left), and quantification of brain temperature during the beginning, middle and end of a torpor bout (right) (n=7, one-way ANOVA with Geisser-Greenhouse correction and Tukey’s multiple comparison, P=0.04 (0 vs. 50%) and P=0.01 (50% vs. 100%)). (D) Histogram showing the relative frequency of EMG events during torpor. (E-G) Average EMG event duration (E), inter-EMG interval (F), and EMG event frequency (G) throughout the torpor bout expressed as percent of torpor bout that has passed (left), and comparison of the beginning, middle and end of a torpor bout (right) (n=7, one-way ANOVA with Geisser-Greenhouse correction and Tukey’s multiple comparison, P=0.01 (EMG duration 0 vs. 50%), P=0.01 (EMG frequency 0% vs. 100%), and P=0.03 (EMG frequency 50% vs. 100%)). (H-J) Correlation between the EMG event duration and frequency (H), EMG event duration and brain temperature (I), and EMG event frequency and brain temperature (J). The average EMG durations and frequencies for each hamster within 5 time bins spanning the torpor bout are plotted (5 time bins from each of the 7 hamsters = 35 data points, individual hamsters are plotted in different colours, Pearson correlation). Error bars are ±SEM.

### Video recordings of hamsters in torpor reveal that EMG events comprise a wide array of behaviours

To determine whether the periods of muscle activity during torpor are due to shivering or other types of behaviour, we performed thermal imaging and video recordings in a separate cohort of three short-photoperiod adapted hamsters undergoing torpor. Torpor bouts were identified from a prolonged drop in skin temperature and the videos were visually inspected to identify behavioural characteristics during periods of movement. Note that thermal imaging experiments were performed in a separate cohort of unimplanted animals, in a separate location at an ambient temperature of 19-21 °C, which is higher than the animals used for EEG and EMG recordings, which were housed at an ambient temperature of 14-18 °C. The average drop in skin temperature in hamsters recorded with thermal cameras was of approximately 7°C and the lowest recorded skin temperature was 23.5 °C. Thermal recordings were used to avoid invasive procedures and disrupt animals as little as possible. Previous work indicates that thermal imaging is a reliable method to detect torpor in a laboratory setting (27). Visual inspection of the entire progression from normothermic behaviours to hypothermia during torpor and back to normothermic body temperatures revealed a wide array of behaviours and showed that movements were regularly observed even during the body temperature nadir (supplementary video 1: ‘50x_Torpor_full’). During torpor, immobility was interspersed with short behavioural events where animals engaged in a variety of behaviours, such as twitching, nesting, repositioning in the nest, scratching/grooming, and even leaving the nest (supplementary video 2: ‘Torpor_movements’). Contrary to our expectation, we did not observe unequivocal periods of shivering, neither during sustained torpor or during arousal from torpor, where behaviours mostly consisted of minor movements, such as twitching, stretching, looking around, and grooming (supplementary video 3: ‘Torpor_rewarming’). In summary, visual inspection of hamsters in torpor at an ambient temperature of 19-21 °C shows that behaviours during torpor EMG events consist of wake-like activities, such as repositioning, nesting, and grooming, while unequivocal periods of shivering were not observed.

### Torpor EMG events are associated with an increase in EEG activity

While torpor in general was characterised by an overall decrease in EEG amplitude, EMG events occurred alongside an increase in EEG power (figure 3A-B). Power spectral density analysis of the frontal EEG signal before and during an EMG event revealed a significant increase in both slow (2-2.5 Hz) activity and across a broad frequency band from 9.5-15.5 Hz (figure 3C). We named this frequency band the ‘movement band’ (M-band). The increase in power during vs. before the EMG event confirmed a large difference within M-band frequencies (figure 3D). Analysis of M-band power before, during, and after an EMG event revealed that the increase in M-band power started ∼30 seconds before the beginning of the muscle activity and peaked ∼6 seconds after muscle activity initiation (figure 3E). On the contrary, EEG slow wave activity (1-4 Hz) decreased prior to the EMG event but then exhibited a sharp increase upon the beginning of muscle activation, followed by a gradual decrease. M-band power amplitude during EMG events correlated positively with both brain temperature (figure 3F) and EMG event duration (figure 3G).

**Figure 3.**
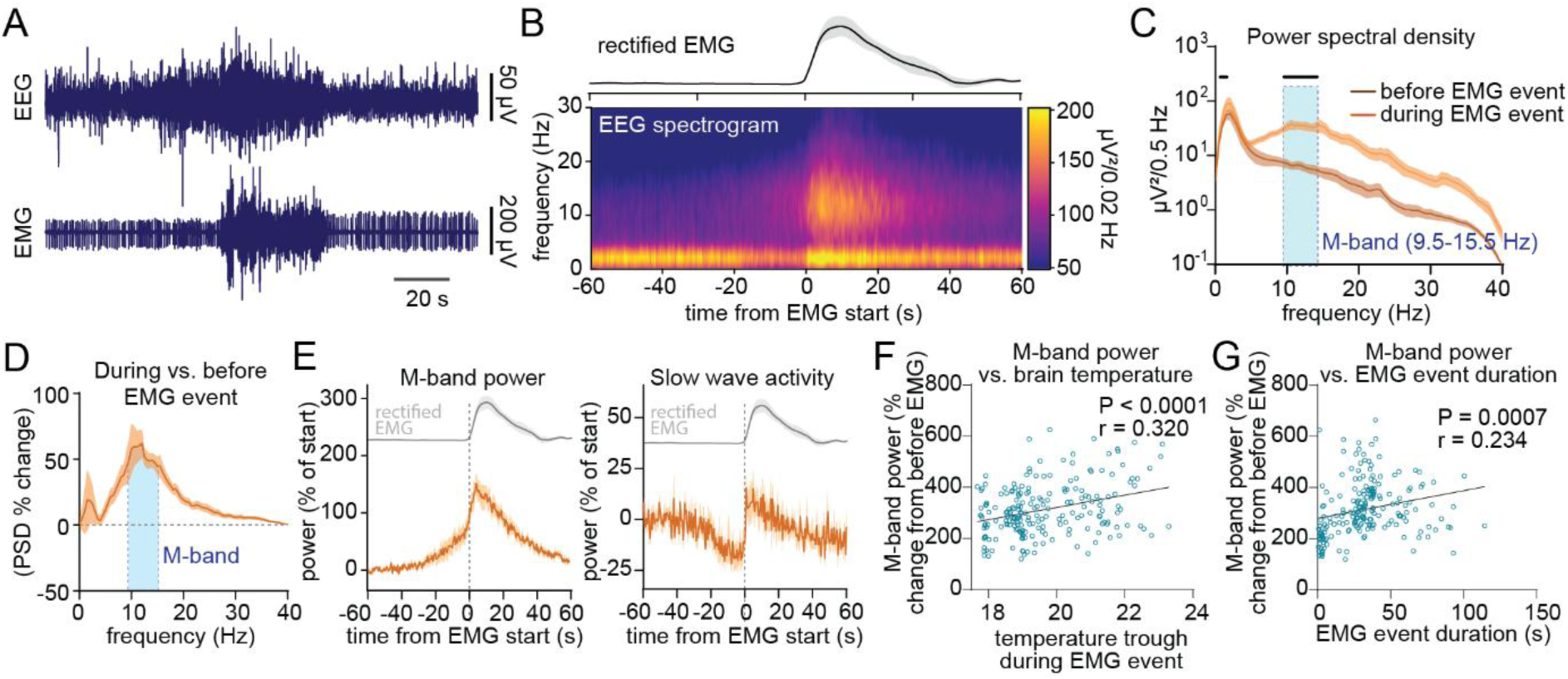
Torpor EMG events are associated with an increase in EEG activity. (A) Example of EMG and EEG trace from a hamster during torpor showing a bout of muscle activity. (B) average rectified EMG signal (top) and spectrogram of the frontal EEG derivation (bottom) time-locked to the start of EMG activity during torpor. (C) Power spectral density analysis of the EEG before (30-60 seconds before muscle activity start) and during (from 0-10 seconds after muscle activity start) (n=7, 2-way ANOVA with Sidak’s multiple comparison, black horizontal lines depict frequencies with significant difference between data points). (D) EEG power during EMG events expressed as percent of the power 30-60 seconds before the EMG event start (n=7). (E) M-band power (9.5-15.5 Hz) (left) and slow wave activity (1-4 Hz) (right) time-locked to the beginning of EMG event and expressed as % of power 50-60 seconds before EMG event start. The top trace shows the average rectified EMG amplitude (n=7). (F-G) Correlation between peak M-band power from data in (E) and minimum brain temperature during EMG (F) and EMG event duration (G) for each EMG event from all hamster (Pearson correlation, 208 EMG events included, 6 outliers removed using ROUT method). Error bars are ±SEM. M-band = movement band.

### Torpor EMG events are often preceded by an ‘evoked-like’ potential

Upon examination of the EEG traces during torpor we noticed that EMG events were often preceded by a distinct EEG pattern resembling an evoked potential consisting of a brief sequence of high-amplitude positive and negative waves. This ‘evoked-like’ potential was immediately followed by an increase in M-band power leading up to the muscle activation 10-30 seconds later (figure 4A). To systematically quantify the concomitant changes in brain and EMG activation, the largest local trough in the EEG signal preceding each EMG event was identified and used as a temporal marker for the evoked-like potential. The analysis showed a clear peak in EEG power during the evoked-like potential, immediately followed by the initiation of a decrease in slow wave activity and increase in M-band power (figure 4B). Importantly, muscle activity did not begin until 10-30 seconds after the evoked-like potential. Quantification of EEG slow wave activity and M-band power before and after the evoked-like potential confirmed that slow wave activity decreased while M-band power increased after its termination (figure 4C-D). Thus, the evoked-like potential delineates the beginning of the M-band rich period leading up to an EMG event.

**Figure 4.**
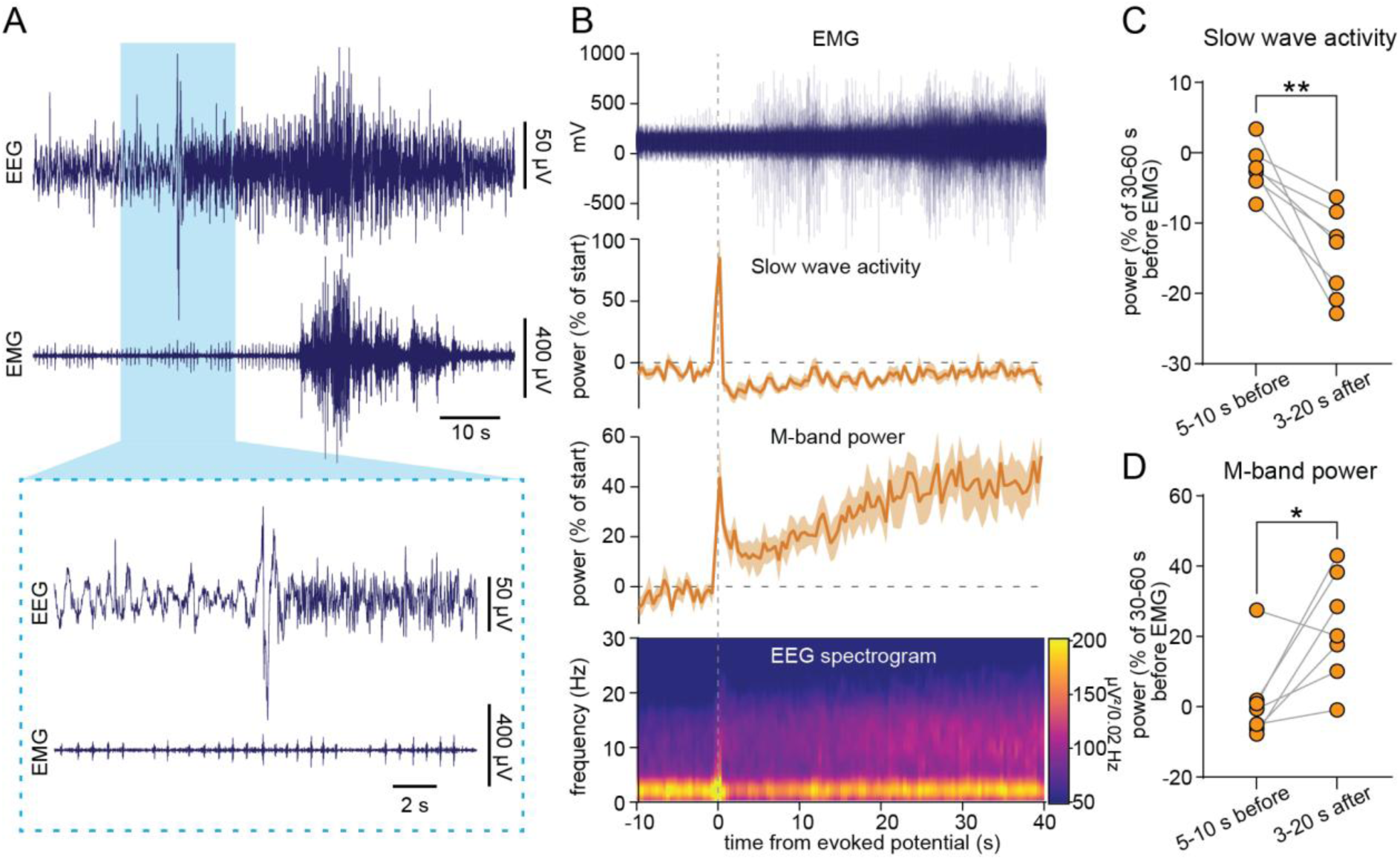
Torpor EMG events are often preceded by an ‘evoked-like’ potential. (A) Example of EEG and EMG trace from a hamster during a torpor EMG event. The insert is higher magnification showing the evoked-like potential and shift in EEG. Note that muscle activity is not initiated until ∼20 seconds after the evoked-like potential. (B) EMG and EEG around the evoked-like potential across all trials. Top: all EMG traces overlayed, middle: slow wave activity and M-band power plotted as the percent of the power 30-60 seconds before the EMG event start, bottom: EEG spectrogram. All are time-locked to evoke-like potentials preceding EMG events (n=7). (C-D) Quantification of the average slow wave activity (C) and M-band power (D) before and after evoke-like potentials that precedes EMG events (n=7, paired t-test, P=0.0042 and P=0.026). Power is presented as percent of the power 30-60 seconds before the EMG event start. Error bars are ±SEM.

### EMG events during NREM sleep and torpor have distinct characteristics

Micro-arousals are brief (typically lasting less than 20s) periods of wake-like brain activity associated with increased muscle tone observed during NREM sleep in both rodents(28–31) and humans (32). We next asked how NREM sleep micro-arousals (NREM EMG events) compare to muscle events during torpor (torpor EMG events). In order to compare NREM EMG events with torpor EMG events in hamsters, we used the same parameters to detect NREM EMG events as we used for torpor EMG events and therefore included all EMG episodes with a duration below 120 seconds, thus encompassing short bouts of arousal, that would conventionally be classified as wakefulness. Analysis of NREM sleep episodes during days without torpor revealed that the hamsters exhibited regular episodes of increased muscle activity during NREM sleep in a similar manner to what has been reported in mice (figure 5A). Most of these NREM EMG events were of a duration between 4-8 seconds long, which corresponds to previous reports on micro-arousals in mice(31) (figure 5B). Comparison of NREM EMG events and torpor EMG events showed that NREM EMG events were significantly shorter on average and occurred with a much higher frequency (torpor EMG events: 30.1 ± 8.2 sec duration, 6.2 ± 2.6 events per hour; NREM EMG events: 15.5 ± 9.5 sec duration, 30.1 ± 8.2 events per hour) (figure 5C-D). NREM EMG events were associated with a decrease in EEG power that was evident across a broad frequency range including slow wave activity (1-4 Hz) and M-band power (figure 5E-G). This decrease in power is comparable to results from mice but is in stark contrast to the relative increase in EEG amplitude during torpor EMG events. Despite the overall relative decrease in EEG power during NREM EMG events, comparison of the power spectral density during torpor and NREM EMG events showed significantly higher frequencies between 1-8 Hz in NREM EMG events compared to torpor EMG events (figure 5H). Comparing the change in power during and before EMG events confirmed that torpor EMG events were associated with a large, relative increase in power while NREM EMG events resulted in a relative decrease in power (figure 5I-J). Finally, NREM EMG events did not exhibit the delayed increase in brain temperature that characterized torpor EMG events. Instead, the temperature peaked during the NREM EMG event and then dropped (figure 5K). Video recordings and thermal imaging of a separate group of hamsters were used to visually assess the characteristics of putative EMG events during immobility under normothermic conditions most likely corresponding to sleep (supplementary video 4: ‘MicroArousals_putative’). The behaviour during periods of movement during sleep consisted mostly of short twitches or minor repositioning in the nest. Contrary to movements during torpor, complex behaviours such as nesting and grooming were not observed. Upon transition from immobility to a longer period of active wakefulness, grooming, nesting and other complex behaviours were present. In summary, EMG activity during NREM sleep in hamsters showed different characteristics from torpor EMG events in terms of duration, frequency, brain activity patterns, brain temperature signatures and behavioural characteristics.

**Figure 5.**
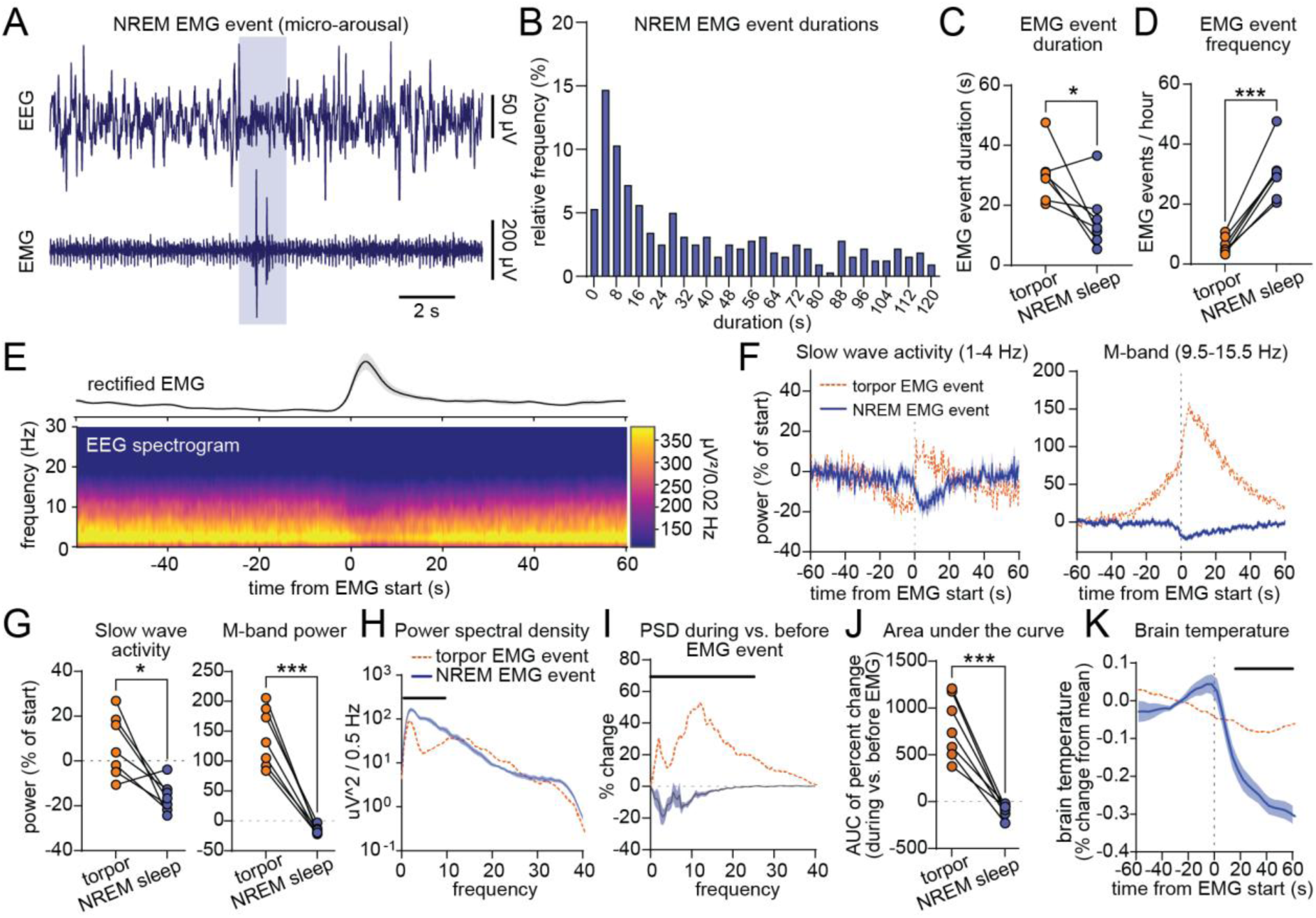
EMG events during NREM sleep and torpor have distinct characteristics. (A) Example of EMG and EEG trace from a hamster during NREM sleep showing a NREM EMG event (micro-arousal). (B) Histogram of the frequency of EMG activity durations during NREM sleep. All EMG activity events with a duration below 120 seconds are included in all analysis. (C) Quantification of average EMG event durations (C) and EMG event frequency (D) during torpor and NREM sleep (n=7, paired t-test, P=0.03 for (C) and P=0.0001 for (D)). (E) Spectrogram of EEG signals time-locked to the beginning of NREM sleep muscle activity. The average rectified EMG signal is shown in the top (n=7). (F) Slow wave activity (1-4 Hz) (left) and M-band power (9.5-15.5 Hz) (right) in blue, time-locked to the beginning of EMG event and expressed as % of power 50-60 seconds before EMG event start. Orange dotted traces show slow wave activity and M-band power during torpor EMG events (n=7). (G) Comparison of slow wave activity (left) and M-band power (right) during torpor EMG events and NREM EMG events (n=7, paired t-test, P=0.02 for slow wave activity and P=0.0001 for M-band power). Power is analysed for 5-10 s after the onset of EMG event, presented as % of 30-60 s before the EMG event. (H) power spectral density plots for torpor and NREM EMG events (n=7, 2-way ANOVA with Sidak’s multiple comparison, black horizontal line denotes significantly different frequencies). (I) comparison of EEG power during EMG events in torpor and NREM sleep expressed as percent of the power 30-60 seconds before the EMG event start (n=7, 2-way ANOVA with Sidak’s multiple comparison, black horizontal line denotes significantly different frequencies). (J) Area under the curve for data shown in (I) (n=7, paired t-test, P=0.0004). (K) Average brain temperature time-locked to muscle activity during NREM sleep and torpor (n=7, 2-way ANOVA with Sidak’s multiple comparison, black horizontal line denotes significant difference). Error bars are ±SEM.

## Discussion

Torpor is typically considered a state of inactivity, lethargy, and low arousal. However, this widely held view is increasingly recognised as overly simplistic, and a closer look at the physiological signals in torpid animals reveals a dynamic state in terms of behaviour, arousal state, brain activity and temperature. In this study we sought to characterise the brain activity and temperature changes that accompany torpor EMG events in the Djungarian hamster to gain a better understanding of their role and implications.

Consistent with previous reports(14, 16, 25), brain temperature, EEG amplitude and EEG power were lower during torpor than euthermic periods of wakefulness, NREM and REM sleep. The correlation between EMG event frequency and absolute brain temperature suggests that the frequency of EMG events is influenced by absolute brain temperature rather than relative temperature changes that occur over the course of torpor. On the contrary, relative brain temperature and EMG event duration exhibited an opposite pattern where the middle of torpor bouts was associated with the lowest relative brain temperatures and longer EMG events. Absolute brain temperature and EMG event duration also showed a significant negative correlation, but weaker than the correlation between brain temperature and EMG frequency. These results indicate that different mechanisms may underlie the regulation of EMG event frequency and EMG event duration. Previous studies(14, 16) have demonstrated that rewarming from torpor is followed by an increase in SWA, which is correlated with the duration and degree of hypothermia and resembles rebound sleep after sleep deprivation. This indicates the existence of a mechanism that keeps track of time spent in torpor. A similar mechanism may be responsible for the increase in EMG event frequency over the course of the torpor bout, since the increase in EMG event frequency did not correspond to changes in brain temperature over the course of the torpor bout. Alternatively, changes in neuromodulatory tone and/or metabolic state may also influence EMG event frequency independent of brain temperature

The apparent link between brain temperature and EMG event frequency and duration, as well as the observation that torpor EMG events were consistently followed by an increase in cortical temperature, point to the possibility that muscle activation may serve a thermoregulatory function during torpor. One possible circuit mechanism that may drive the generation of movement in response to low temperature could involve the hypothalamus and spinal pathways. During torpor, cooling of the brain could be detected by temperature sensing areas such as the preoptic area, which would disinhibit the dorsomedial hypothalamus(24, 35). The dorsomedial hypothalamus could then activate the rostral medullary raphe neurons, which drive spinal motor neurons to produce EMG bursts and stimulate sympathetic thermogenesis in brown adipose tissue, as what has been shown previously in rats(36, 37). These EMG events would then generate heat and, along with increased blood flow, transiently raise cortical temperature(37, 38). Arousal systems and sleep-pressure signals could potentially play a role in modulating the frequency and duration of these EMG events during torpor(39, 40). An intriguing question is what controls the initiation of the EMG events. One possibility is that EMG events are initiated by a mismatch between the setpoint for the lowest ‘allowed’ temperature and the actual temperature, although the exact brain areas responsible for this are yet to be defined. Such a setpoint may be absolute or be dynamically regulated over the course of the torpor bout. Opposed to the small increase in temperature brought upon by single EMG events, arousal from torpor is associated with a steep increase in temperature(41). We cannot deduce from this study whether torpor EMG events and arousal from torpor are initiated by the same physiological mechanisms or if they are controlled by separate systems.

Previous reports on hibernation(19) or fasting-induced torpor(33) have shown that these hypometabolic states are punctuated with shorter periods of wake-like activity that are usually accompanied by increases in body temperature. During these periods, animals have been observed to shiver, groom or build nests. In the current study, video recordings and concurrent thermal imaging of hamsters in torpor support that the behaviour performed during torpor EMG events are more similar to behaviours seen during wakefulness, including grooming and locomotion but opposed to previous reports we did not observe shivering in torpid, uninterrupted hamsters. This may indicate that EMG-events observed here are more likely to be associated with non-shivering thermogenesis. It is important to note that the hamsters used for visual inspection of torpor EMG event behaviours were housed at a higher ambient temperature (19-21 °C) than the hamsters used for analysis of EEG, EMG and brain temperature, which were housed at an ambient temperature of 14-18 °C. A low ambient temperature is not required for entry to torpor in Djungarian hamsters that are already prone to torpor entry, and torpor characteristics, such as torpor bout duration, is not dependent on the ambient temperature(34). Still, we cannot rule out that shivering may be more evident under lower ambient temperatures. Therefore, future studies are needed to confirm if EMG events under lower temperature also includes shivering.

An interesting feature of the EMG events is that they were often preceded by an ‘evoked’ like potential (thus called due to its large, positive and negative peaks characteristic of evoked potentials), which was followed by a change in the EEG to faster frequencies within a broad range of 9.5-15.5 Hz. We dubbed this frequency band, where the difference was most prominent, the ‘movement band’ or M-band in short. Figure 6 shows a graphical summary of the specific EEG patterns associated with an EMG event, including the evoked-like potential and the increase in M-band frequencies prior to the muscle activity. An important question is whether the evoked-like potential before the shift in EEG has a function or if it merely reflects a change in arousal state. Since the recordings in this study were conducted under tightly controlled conditions, this evoked-like potential is unlikely to occur in response to external stimulation. Therefore, we hypothesise that it may reflect an intrinsically evoked potential. A type of intrinsically evoked brain potential previously described is the heartbeat evoked potential (HEP)(42–45). HEPs have been reported in a number of human studies on interoceptive processing and also in recent work in rats, where increasing or decreasing the heart rate using cardiac muscle stimulation and vagus nerve stimulation, respectively, led to the appearance of distinct cortical and hippocampal EEG and LFP characteristics. Interestingly, the movement and electrical signals of the gut and stomach have also been linked directly to brain activity during sleep(46, 47), and stimulation of the abdomen or intraperitoneal tissue can induce evoked responses in the brain EEG(48). This relationship between body and brain inspired the visceral hypothesis of sleep, which states that the brain is primarily engaged in processing of external information during wakefulness, but switches to processing of interoceptive information during sleep(49). Less is known about the interoceptive processing during torpor, but our results suggest that EMG events during torpor may result from interoception rather than external stimuli. Indeed, torpor occurs alongside pronounced changes in cardiovascular, respiratory and visceral rhythms, which may result in a more salient interoceptive signaling.

**Figure 6.**
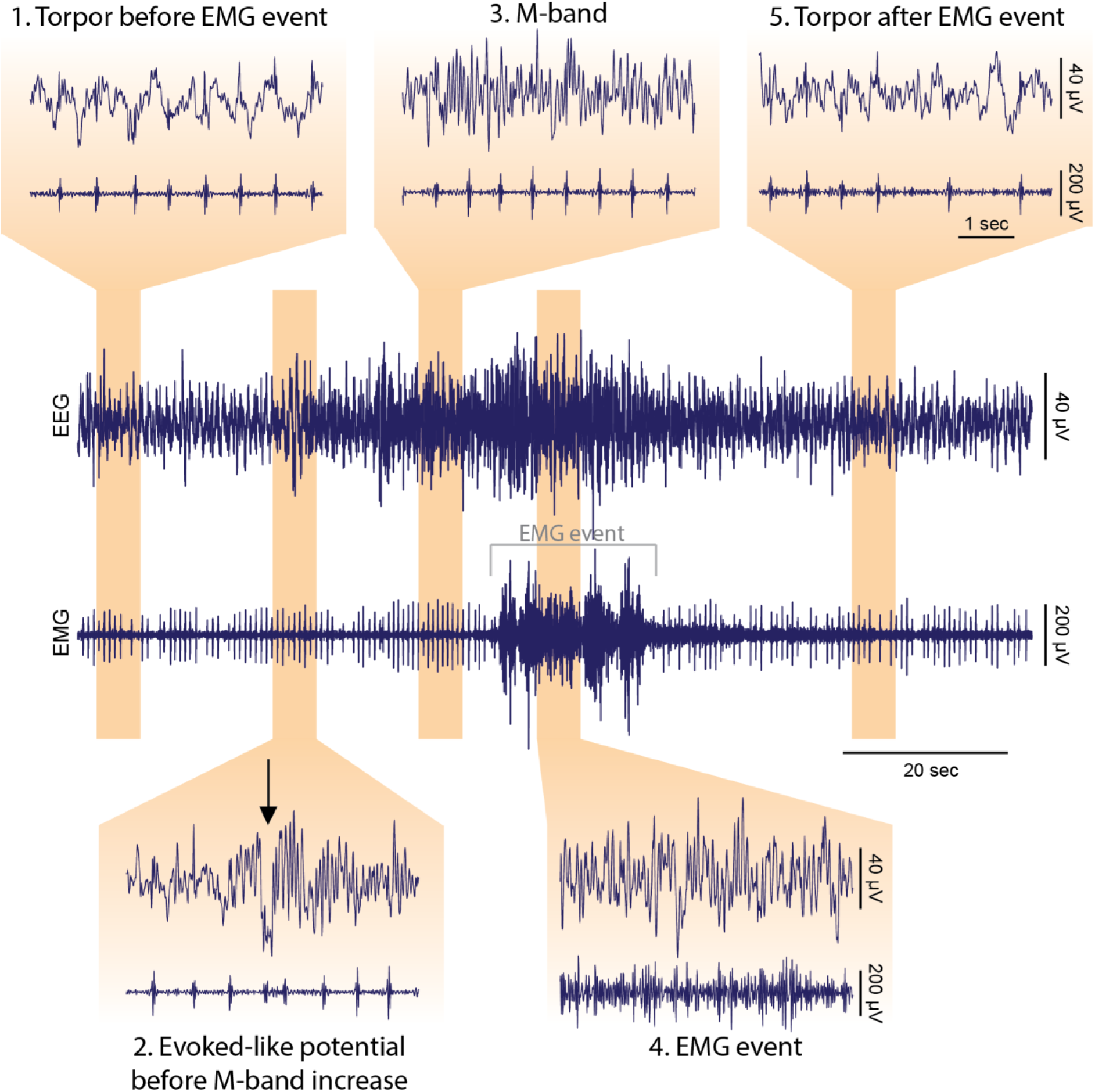
Example traces showing the different phases of a torpor EMG event. (A) EEG and EMG signals during one EMG event during torpor. Each orange box depicts separate time periods before, during or after the EMG event. (1) The EEG activity during torpor is characterized by low amplitude and there is a lack of EMG tone. (2) An evoked potential delineates the transition from typical torpor EEG with increasing power in the M-band (9,5-15.5 Hz). (3) The period between the evoked potential and the EMG event is characterized by high M-band power but no EMG activity. (4) During the EMG event, the EEG M-band power is peaking and EMG activity is high. (5) After the EMG event, muscle activity stops and the EEG returns to a low amplitude level similar to before the evoked potential.

Our analysis showed that torpor EMG events and NREM micro-arousals have different characteristics and are associated with distinct effects on the brain and body. Torpor EMG events are longer in duration and occur less frequently than NREM micro-arousals. During NREM micro-arousals, EEG SWA and M-band power decrease, though power in the 1–8 Hz range remains higher than during torpor EMG events. In contrast, torpor EMG events are associated with increases in overall EEG power. Furthermore, while NREM micro-arousals coincide with decreasing brain temperature, torpor EMG events are consistently followed by an increase in brain temperature. We therefore propose that torpor EMG events are driven primarily by thermoregulatory pathways and that their primary function is thermogenic corrections. NREM micro-arousals are thought to arise from transient activation of arousal systems, including regular noradrenergic bursts from the locus coeruleus (LC) at an infraslow timescale of around 0.02 Hz (30, 50). The infraslow noradrenaline oscillations during NREM sleep are implicated in memory consolidation(30), sleep state transitions(51), and brain clearance via the glymphatic system(52). It is to be established whether infraslow noradrenaline dynamics also exist during torpor and if LC activity has a role in controlling torpor EMG events. This is an intriguing question as it would point to whether the properties such as brain clearance take place during torpor. The infraslow rhythm is conserved across amniotes, including lizards, rats, mice, humans, and pigeons, indicating that it may be a fundamental feature in the evolution of vigilance states(53, 54). The existence of periodic motility in torpor suggests that infraslow dynamics, albeit on a slower time scale than what is observed during normothermic NREM sleep in rodents, extends to states of dormancy, further supporting the notion that infraslow dynamics are of high evolutionary importance.

In conclusion, our analysis showed that EMG events, associated with an increase in brain temperature and EEG power of slow and fast EEG frequencies are an inherent phenomenon of naturalistic daily torpor in Djungarian hamsters. Although the underlying mechanism driving these EMG events is still unclear, the presence of an evoked-like potential in the EEG prior to the occurrence of EMG events may reflect an internal sensory process, which potentially may protect the animal from excessive cooling. Finally, the difference between torpor EMG events and NREM micro-arousals indicate that these events may have distinct functions, although future studies are needed to investigate potential similarities and differences in the circuitry underlying the two events.

## Author contributions

T.D. and R.Mu. performed the experiments, N.L.H. and R.Mu. performed the analysis and made the figures, X.Z., A.H-S., R.Ma., S.N.P., A.H., T.D. and V.V.V. provided technical support, advice and project management, N.L.H., R.Mu., T.D., and V.V.V wrote the manuscript, all authors edited the manuscript.

## Acknowledgements

We thank Dr S. Palchykova for collecting the original data used for EEG/EMG analysis, Prof. Irene Tobler for detailed comments and feedback on the manuscript, Simona Di Pretoro for help with animal husbandry and colony maintenance, and all members of VV lab as well as Biomedical Services staff for help with animal husbandry and assistance with experiments.

## Supplementary material

Supplementary videos are uploaded to Zenodo (https://doi.org/10.5281/zenodo.18363686) under DOI: 10.5281/zenodo.18363685.

## Methods

### Part I: Electrophysiology and brain temperature recordings

#### Animals

The data was collected at the University of Zurich in 2001 and used in a previous study(55). 7 Adult Djungarian hamsters (*Phodopus sungorus*) (eight males) were used. Hamsters were housed individually in Macrolon cages (36 x 20 x 35 cm) with food and water available *ad libitum*, and maintained in an 8 h light-16 h dark cycle [light from 09:00 to 17:00 h; 7 W OSRAM DULUX EL energy saving lamp (Osram, Germany), approximately 30 lux], and a mean ambient temperature of 14-18 °C. Rest-activity behaviour was monitored continuously with IR-sensors, and weight and pelage colour were scored every week. After 10 weeks, animals showing pronounced adaptation to the short photoperiod based on rest-activity behaviour, decreased body weight, and an increase in fur colour index, were implanted with EEG and EMG recording electrodes and thermistors to record brain temperature as well as a temperature sensitive transmitter implanted I.P. to continuously record body temperature (Mini Mitter) (body temperature data was not analysed in this work).

#### Surgery

Hamsters were anaesthetized (Ketalar, 75 mg/kg, Parke-Davis, Pfizer AG, Zurich, Switzerland; Rompun 4 mg/kg, Bayer, Leverkusen, Germany) and implanted with gold-plated miniature screws (0.9 mm diameter) inserted into the skull to serve as EEG electrodes. Screws were placed epidurally over the right frontal cortex (2–3 mm lateral to midline and 2 mm anterior to bregma) and right parietal cortex (2 mm lateral to midline and 2 mm posterior to bregma). In all animals a reference electrode was placed over the cerebellum (2 mm posterior to lambda, on midline). A calibrated thermistor (Thermometrics, Inc., Edison, NJ, USA, P20, R 25 °C at 1 kW, maximum diameter 0.5 mm, accuracy ±0.05 °C) was inserted between the skull and dura through a hole over the left frontal cortex (2–3 mm lateral to midline and 2 mm anterior to bregma) to record cortical brain temperature. Two gold wires were inserted into the neck muscles (diameter 0.2 mm) to record the electromyogram (EMG). The electrodes and thermistor were connected to stainless steel wires that were fixed to the skull with dental cement. At least 1 week was allowed for recovery before the recording was started.

#### Signal acquisition and vigilance state scoring

EEG and EMG signals were recorded continuously over several days. The EEG and the EMG signals were amplified (amplification factor approx. 2,000), conditioned by analog filters (high-pass filter: −3 dB at 0.016 Hz; low-pass filter: −3 dB at 40 Hz, less than −35 dB at 128 Hz) sampled with 512 Hz (digitally filtered, EEG: low-pass FIR filter 25 Hz; EMG: band-pass FIR filter 20-50 Hz) and stored with a resolution of 128 Hz. Sleep stages were scored visually based on the EEG and EMG signals in 4-second epochs. Waking was characterized by low voltage, high-frequency EEG pattern and phasic EMG activity. NREM sleep was characterized by the occurrence of high amplitude slow waves and low tonic EMG activity. During REM sleep the EEG was similar to that during waking, but only heart beats and occasional twitches were evident in the EMG signal. Thermistor data was recorded every 4 seconds.

#### Experimental design

For each hamster, a day with no attempt to enter torpor served as baseline and a day with torpor served as torpor day. If a hamster showed multiple days of torpor, one of the days were selected for further analysis based on visual inspection of signal quality and torpor duration. Analysis of baseline day data was performed from Zt1 to Zt12. Analysis of torpor was performed when the cortical temperature was within 10% of the lowest temperature recorded during the torpor bout. Division of torpor bouts into bins based on the percent time of the torpor bout that has passed was done by adding the total duration of a torpor bout (defined as cortical temperature within 10% of lowest recorded temperature) and dividing it into 5 bins equivalent of 0-25%, 25-50%, 50-75%, and 75-100% of the torpor bout (average bin duration = 58.2±8.5 minutes).

#### Automatic muscle activity detection

The detection of muscle activity was based on the EMG signal and performed in MATLAB. For muscle activity detection during torpor, only muscle activity occurring when the brain temperature was within 10% of the lowest recorded temperature was included. Periods scored as wakefulness or REM sleep during torpor were re-scored as torpor if the duration of the period was below 60 seconds in order to remove existing manually scored vigilance states. The EMG signal was then filtered and rectified using the moving standard deviation, the moving mean and a 10000^th^ order median filter. Peaks in the EMG signal were detected and stored if their magnitude crossed a threshold that was selected for each individual hamster based on visual inspection of the data and the automatically scored peaks in muscle activity. In the subsequent steps, EMG peaks were excluded if they did not occur within torpor, and EMG peaks that were less than 10 seconds apart were combined as one event.

For detection of muscle activity during NREM sleep, the same parameters were used but new thresholds were selected for each hamster and NREM sleep during a day without entry to torpor was used. For one hamster, the algorithm was not able to properly detect peaks in muscle activity due to high amplitude heart beat artifacts so for this hamster, micro-arousals were selected based on the manual vigilance scoring where wake periods with a duration below 120 seconds during NREM sleep were stored as micro-arousals.

#### Data analysis

All data analysis was performed in MATLAB (R2024a). Prior to spectral analysis, data traces were filtered using a Cheby1 filter with stop band frequencies between 0.5 and 64 Hz and Passband frequencies between 1 and 50 Hz. The lower passband of 1 Hz was chosen to exclude potential low frequency artefacts caused by movement during muscle activity.

##### Spectrograms

Spectrograms were generated using the ‘spectrogram’ function with a window of 500 ms and an overlap of 50 ms. Mean power within specific power bands was calculated from the spectrogram by averaging the power within the frequencies of interest. Power spectral density (PSD) plots were calculated using pwelch with a frequency resolution of 0.5 Hz, zero overlap and a Hanning window. PSD plots from before, during and after a muscle activity were calculated 30-50 seconds before the muscle activity, 0-2 seconds after the beginning of the muscle activity and 5-10 seconds after the beginning of the muscle activity, respectively.

##### Calculation of average rectified EMG

Average rectified EMG traces were calculated by taking the mean of the filtered and rectified EMG trace time-locked to each muscle activity.

#### Statistics

Statistics were done in GraphPad Prism (version 10.3.1). P-values less than 0.05 (after correction for multiple comparison except the comparison of power density per frequency bins) were considered significant. For comparison of the means of two group where the groups consisted of the same hamsters under different conditions or time bins, a two tailed paired t-test were used. For comparison between the means of three or more groups, RM one-way ANOVA with Geisser-Greenhouse correction and Tukey’s multiple comparison was used. For comparison of power densities across frequency bins RM two-way ANOVA with matched values stacked into sub columns was used. Pearson correlation was used to calculate correlation coefficients and slopes of the correlations. Outliers were detected using the ROUT method and any removed data points are mentioned in the figure legend.

### Part II: Video and thermal imaging recordings

#### Animals

The data was collected at the University of Oxford in 2025. Three adult (6-8 months old) Djungarian hamsters (*Phodopus sungorus*) (all females) were used. Hamsters were singly housed in (Tecniplast blue-line IVC cages used as open-tops) with food and water available *ad libitum*, and maintained in an 8 h light-16 h dark cycle [light from 09:00 to 17:00 h; 2.5 W, 6000°K Campden Instruments LED Light Bar, 488.18 photopic lux, S-cone-opic lux: 14.99, Melanopsin-opic lux: 406.09; Rod-opic lux: 427.39; M-cone-opic lux: 437.70 – measured by nanolambda digital spectrometer], and a mean ambient temperature of 19-21 °C. The ambient temperature of hamsters used for video recordings was higher than the ambient temperature of the animals used for EEG/EMG and brain temperature recordings due to inaccessibility to facilities where the temperature could be reduced further. The weight and pelage colour of all hamsters were scored every week. After 10 weeks, animals showing pronounced adaptation to the short photoperiod based on decreased body weight, and an increase in fur colour index, were moved into recording cages under the same light cycle and ambient temperature (custom-built open-top plexiglass cages: 24.4 cm (W) x 34 cm (D) x 39 cm (H), corresponding to ca. 833 square cm floor space). These recording cages were housed inside sound-attenuated recording chambers [light from 09:00 to 17:00 h; 2.5 W, 6000°K Campden Instruments LED Light Bar, 488.18 photopic lux, S-cone-opic lux: 14.99, Melanopsin-opic lux: 406.09; Rod-opic lux: 427.39; M-cone-opic lux: 437.70 – measured by nanolambda digital spectrometer]. Here, their skin/fur temperature was recorded non-invasively with thermal imaging cameras (Optris Xi 80 compact spot finder thermal imaging camera with 80° wide angle lens, Optris GmbH, Berlin, Germany; 1Hz sampling frequency). The pixel reporting the hottest temperature was used as the skin/fur temperature. Infra-red video cameras (ELP USB 1080P HD Camera with IR LED, acquired at 30 frames per second) were also mounted on top of the recording cage on custom-made 3D-printed camera holders. The recordings were made for 2-3 weeks.

#### Thermal and video data acquisition and processing

The animals’ skin/fur temperature and ambient temperature was calculated as the hottest pixel in view and a 2*2-pixel spot with the lowest mean temperature, respectively. This data was saved every second. The behavioural video was recorded at a frame rate of 30 frame per second. While all video cameras had IR night vision, day vision was either colour or black and white. All cameras were restarted daily at dark onset.

Thermal imaging data was then processed by computing a moving average of the temperature data to smoothen abrupt, non-physiological changes in temperature due to movement of the animals. This thermal imaging pipeline had previously been validated against intra-abdominal temperature sensors(27).

An animated plot of skin/fur temperature of each day was created in MATLAB and combined with the behavioural video in CapCut (Desktop Version 7.3.0, https://www.capcut.com/). All video editing was done using MATLAB and CapCut.

